# Seasonal and genetic effects on lipid profiles of juvenile Atlantic salmon

**DOI:** 10.1101/2023.02.22.529528

**Authors:** Andrew H. House, Paul V. Debes, Minna Holopainen, Reijo Käkelä, Iikki Donner, Morgane Frapin, Ehsan Pashay, Johanna Kurko, Hanna Ruhanen, Craig R. Primmer

**Author notes:** **Corresponding Author:** Craig Primmer, **Address:** Viikinkaari 9 (PL 56), 00790 Helsinki, Finland.

## Abstract

Seasonality can influence many physiological traits requiring optimal energetic capacity for life-history stage transitions. In Atlantic salmon, high-energy status is essential for the initiation of maturation. Atlantic salmon lipid reserves are predominantly found in the viscera and myosepta in the muscle while the liver is essential for maintaining lipid metabolism. A genomic study found a region including a transcription co-factor-coding gene, *vgll3*, linked to Atlantic salmon maturation timing, which acts as an inhibitor of adipogenesis in mice, and mediates maturation via condition factor in Atlantic salmon. Here we investigate the influence of season and *vgll3* genotypes associating with early (EE) and late (LL) maturation on lipid profiles in the muscle and liver in juvenile Atlantic salmon. We reared Atlantic salmon for two years until the occurrence of sexually mature males and sampled muscle and liver at two time points: spring and autumn of the second year. We found no seasonal or genotype effect in lipid profiles in muscle of immature males and females. However, in the liver we did detect a triacylglycerol (TG) enrichment and a genotype specific direction of change in membrane lipids, phosphatidylcholine (PC) and phosphatidylethanolamine (PE), from spring to autumn. Specifically, from spring to autumn membrane lipid concentrations increased in vgll3*EE individuals and decreased in vgll3*LL individuals. This could be explained with two possible scenarios 1) a seasonally more stable capacity of endoplasmic reticulum (ER) functions in *vgll3*\*EE individuals compared to *vgll3*\*LL individuals or 2) *vgll3*\*LL individuals storing larger lipid droplets from spring to autumn in the liver compared to *vgll3*\*EE individuals at the expense of ER capacity. This genotype specific seasonal direction of change in membrane lipid concentrations provides more indirect evidence that a mechanism linking *vgll3* with lipid metabolism and storage exists.

**Highlights:**

- Seasonal lipid species profile separation in muscle and liver in juvenile Atlantic salmon
- Genotype specific direction of change of membrane lipids from spring to autumn
- Indirect evidence that a mechanism linking *vgll3* with lipid metabolism and storage exists

## Introduction

Many physiological traits of fish change in a seasonal manner. The appropriate timing of such changes is critical for the optimal transition between life-history stages, such as the transition from an immature to a mature individual, which require sufficient energy reserves, such as lipids, to be achieved successfully (N. Jonsson & Jonsson, 2003; Rowe et al., 1991; Taranger et al. 2010). Lipid metabolism involves synthesis of membrane phospholipids for physiological functional capacity and synthesis of triacylglycerol (TG) to be able to fuel the metabolic functions. Thus studying lipid metabolism helps to understand how life-history progressions are achieved (reviewed in Tocher, 2003). Due to its predominantly anadromous life-history strategy, Atlantic salmon must rapidly transition between energy usage and storage in order to achieve the necessary physiological changes required for sexual maturation at varying ages and sizes (B. Jonsson & Jonsson, 2005; Post & Parkinson, 2001). Historically, body condition (the relative mass of an individual given its length), also referred to as condition factor, has been used as a proxy of an individual’s lipid reserves and thereby, its energy status (Herbinger & Friars, 1991; Schulte-Hostedde et al., 2005; Sutton et al., 2000). Direct quantification of body lipid reserves has also been used to study the role of lipids in the maturation of fish (Shearer & Swanson, 2000). Currently, detailed mass spectrometric profiling of structurally diverse lipid species enables acquiring an even more accurate view on an individual’s physiological status (reviewed in Rey et al., 2022).

Atlantic salmon is a species accumulating high lipid content in various tissues (Henriques et al., 2014; Vuorinen et al., 2020). Looking at seasonal metabolic changes of salmon juveniles, and identifying tissue specific roles for lipids, may give insight into how metabolic differences of the juveniles influences their energy reserves, growth and maturation. The fatty acids are mobilized from the main triacylglycerol (TG) storage locations, which are in the myosepta in between muscle fibers and the visceral cavity along the intestine (Henderson & Tocher, 1987; N. Jonsson et al., 1997; Morgan et al., 2002; Sheridan, 1988). TG is the primary long-term energy source in most aerobic organisms (Yeo & Parrish, 2022), while membrane lipids, largely comprised of phosphatidylcholine (PC) and phosphatidylethanolamine (PE), respond to thermal acclimatization and contribute to maintaining activities of integral proteins (Kraffe et al., 2007). If TG reserves become limited, also the membrane lipids can be broken down for energy, but at the cost of physiological performance (Tonning et al., 2021). The liver is an important organ for fatty acid and lipid synthesis and lipoprotein production but also serves as the main regulator of lipid metabolism (Jensen-Urstad & Semenkovich, 2012; N. Jonsson et al., 1997; Sissener et al., 2017; Yeo & Parrish, 2022). Mobilizing lipids is key to enabling transitions between major life stages and ultimately survival (Manor et al., 2014; Tocher, 2003). Hence, alterations in the liver and muscle contents of storage and membrane lipid profiles can help us understand differences in physiological traits between individuals.

Sexual maturation is a process that involves massive energy investments and depletions and the age at which this occurs can have dramatic effects on individual fitness (N. Jonsson et al., 1991). Partly for these reasons, understanding environmental, genetic and physiological factors affecting age at maturity in Atlantic salmon has been a popular research topic for decades. In addition, salmon age at maturity is also of applied importance in aquaculture, natural population conservation and sustainable management (reviewed in Mobley et al., 2021). A genome-wide association study earlier identified a genome region including a transcription co-factor-coding gene, *vgll3 (vestigial-like family member 3*), found to be linked to Atlantic salmon maturation age with two alleles, E and L, associating with earlier or later maturation, respectively in male and female salmon (Ayllon et al., 2015; Barson et al., 2015). *Vgll3* had been found to inhibit adipogenesis in mice, (Halperin et al., 2013), and adiposity is well known to promote puberty in many fishes, including salmon (Taranger et al. 2010), thus making *vgll3* effects on adiposity a plausible mechanism influencing age at maturity in salmon. The association between *vgll3* and salmon maturation timing has been supported in several common garden experiments (Åsheim et al., 2023; Ayllon et al., 2019; Debes et al., 2021; House et al., 2021; Sinclair-Waters, Nome, et al., 2022; Sinclair-Waters, Piavchenko, et al., 2022). Further supporting an association with body energy allocation, several studies have identified body condition differences between individuals with alternative *vgll3* genotypes, *vgll3*EE* juveniles having higher body condition (Debes et al., 2021) and more stable body condition throughout the year (House et al., 2023). This stability, in particular the maintenance of higher body condition in the spring, was suggested to contribute to the earlier maturaiton of males with the *vgll3*\*EE genotype in autumn (House et al. 2023). Body condition, however, only provides a very rough approximation of body lipid reserves, and cannot provide information about the relative role of different lipid classes in different tissues. Therefore, a more detailed assessment of lipid classes and individual lipid species profiles within each lipid class is warranted in order to better understand the metabolic processes and capabilities during salmon life history stage transitions.

Here we address this knowledge gap by investigating tissue specific TG, PC and PE lipid classes, and lipid species profiles within each class, in the context of seasonality and *vgll3* genotype in juvenile male and female Atlantic salmon. We reared salmon juveniles from fertilization for two years and assessed temporal, environmental and genetic *vgll3* effects, as well as their interactions, on lipid profiles in the muscle and liver.

## Methods

### Salmon material and sampling

Atlantic salmon juveniles used in this study were the offspring of a first-generation Atlantic salmon hatchery stock maintained by the Natural Resources Institute Finland (62°24’50’’N, 025°57’15’?, Laukaa, Finland). In October 2017, unrelated adults with homozygous *vgll3* genotypes were crossed to create 24 families (six 2 x 2 factorials) where each factorial included a *vgll3*EE* male and female and a *vgll3*LL* male and female. Eggs of each family were divided and incubated in two replicate vertical incubators at ~4.78 °C before transfer to Lammi Biological Station (61°04’45”N, 025°00’40’’E, Lammi, Finland) on April 28^th^ 2018, when they approached the developmental age of first feeding (alevin). Roughly equal numbers of individuals from each family were then placed in five flow-through circular tanks (diameter 90 cm) with water sourced from a nearby lake, Lake Pääjärvi, following the natural annual water temperature cycle (Figure 1). The temperature range for the individuals during the course of the experiment was 1.30-19.04 °C. The average water temperature individuals experienced across the entire experiment was 9.14 °C. Fish were fed *ad libitum* for the duration of the experiment with commercial fish food, the pellet size of which matched the requirements set by the size distribution of the individuals (Raisio Baltic Blend; Raisio Oy). We euthanized and collected tissues from initially 145 individuals (we used less for lipid analyses) at two different time points during this experiment, May and October 2019, representing different seasons, and thus will be referred to as spring and autumn respectively. Liver and muscle tissue for lipid analyses were weighed and flash frozen while gill tissue for determination of smoltification status (see below) was placed in RNAlater for later laboratory analysis. At both sampling periods, individuals were fasted for 24 h and then euthanized by anesthetic overdose of tricaine methanesulphonate (sodium bicarbonate-buffered). Wet mass (± 0.01 g) and fork length (± 1 mm) were measured and gonad development checked to determine maturity status as described in Debes et al. (2021) and a fin clip sampled for genetic analyses. Samples were genotyped with 141 single nucleotide polymorphisms (SNPs) and a sexing marker (Aykanat et al., 2016), and the genetic information used to subsequently determine the *vgll3* genotype and sex of each individual, as well as to assign them to their family of origin as outlined in Debes et al. (2021).

**Figure 1:**
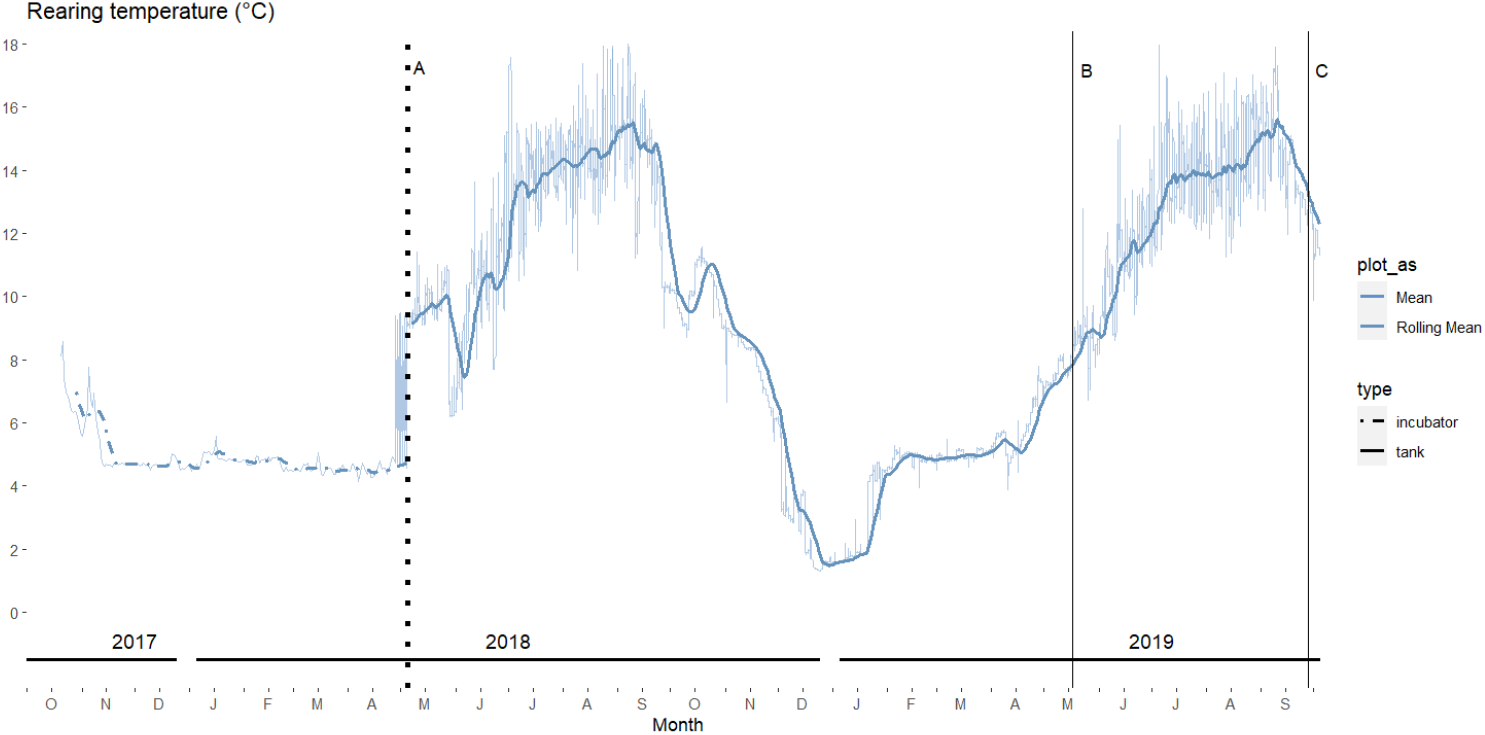
Temperature curve for the experiment from fertilization to final sampling time point. **A)** Transfer date to Lammi Biological Station; **B)** and **C)** are spring and autumn sampling time points, respectively, with routine measurements and tissue sampling of 145 individuals

### Smolt ID Gene expression

A proportion of individuals were observed to have undergone the smoltification process (a physiological and behavioral transition enabling migration from fresh to salt water) in the spring when the first samples were taken. As smoltification can affect lipid storage and use (Sheridan, 1989), and thus potentially confound detection of changes in lipid profiles related to maturation, we limited our study on individuals lacking clear morphological and gene transcription signs and thus indicating a commencement lack of the smoltification process. Transcriptional signs were assessing by reverse-transcription quantitative PCR (RT-qPCR) using the RNA ratio of two gill-expressed marker genes, *atp1a1a.1α* and *atp1a1a.1β*, that have earlier been used for identification of smoltification in Atlantic salmon, and for two already validated stable reference genes in gills; *dnaja2a* and *ef1a* (Piironen et al., 2013). Gills stored in RNAlater were homogenized prior to extraction with a bead mill homogenizer, Bead Ruptor Elite (Omni International Inc.), and RNA was isolated using the NucleoSpin RNA kit (Macherey-Nagel GmbH & Co. KG) and assessed with NanoDrop ND-1000. cDNA synthesis was performed for 57 samples using 500 ng of RNA per sample and the iScript cDNA Synthesis Kit (Bio-Rad Laboratories, Inc.). qPCR primers for the abovementioned genes were designed as described in Ahi & Sefc, (2018) using the online tools OligoAnalyzer 3.1 (Integrated DNA Technology) and Primer Express 3.0 (Applied Biosystems, CA, USA). Primer efficiencies (E) were calculated through standard curves of serial dilutions of pooled cDNA (random samples) and the following formula: E = 10 [−1/slope] (Supplementary data Table S1). The qPCR reactions were prepared as described in Ahi & Sefc, (2018), using PowerUp SYBR Green Master Mix (Thermo Fischer Scientific), and the Bio-Rad CFX96 Touch Real Time PCR Detection system (Bio-Rad, Hercules, CA, USA). To detect the smoltification status, the ratios of the RQ values of the two marker genes (RQ of *atp1a1a.1α* / RQ of *atp1a1a.1β*) were determined as described by Piironen et al., (2013). Fish with a relative expression ratio < 1 were considered as having commenced smoltification (n = 114), and were not considered for further analysis.

### Sample Selection and Lipid Analysis

Samples were selected with the aim of having similar numbers per time point, *vgll3* genotype, and sex for both tissues with final sample sizes being n = 30 for liver, and n = 38 for muscle. Muscle and liver were thawed and a piece (4.0-141.8 mg) was cut and weighed before lipid extraction according to the chloroform and methanol based protocol of Folch et al., (1957). A standardized amount of each lipid extract was diluted in chloroform/methanol 1:2 for a volume of 5 μl then injected into the Agilent 1290 Infinity HPLC system. Chromatographic separation was conducted in a gradient mode with a Luna Omega C18 100 Å (50 x 2.1 mm, 1.6 μm) column (Phenomenex), and employing an acetonitrile/water/isopropanol-based solvent system (Breitkopf et al., 2017) with the flow rate of 0.200 ml/min and 25 °C as the column temperature. Internal lipid standards TG 14:0/14:0/14:0 (NU-CHEK PREP), PC 14:1/14:1 and PE 14:0/14:0 (Avanti Polar Lipids) were included with each sample. The column eluent was infused into the electrospray source of an Agilent 6490 Triple Quad LC/MS with iFunnel Technology and spectra were recorded using both positive and negative ionization modes. TG species were detected as [M+NH4]^+^ ions from MS+ scan. PC species were identified from a Precursor ion 184 scan and quantified from a MS+ scan. Additionally, PE species were identified from a Neutral loss 141 scan and quantified from a MS-scan. Spectra were extracted from the chromatogram with Agilent MassHunter Qualitative Navigator v B.08.00 according to known elution time windows for TG, PC, and PE, and the individual lipid species in each class were identified and quantified using LIMSA software on Excel according to Haimi et al. (2006). Lipid species below 0.5 mol% in a given tissue were removed from the analysis. Concentration values were calculated as pmol/mg tissue. The lipid (isobaric) species are marked as follows: [sum of acyl chain carbons]:[sum of acyl chain double bonds] (e.g., 38:4). Total concentration of each lipid class was calculated by summing up all lipid species concentrations of a class for each individual. For each class of lipid, the species concentration data were used to calculate mol% species profile. Neutral lipid (TG) versus membrane lipid (PC & PE) ratios are also reported in Table 1 and Table 2.

**Table 1:**
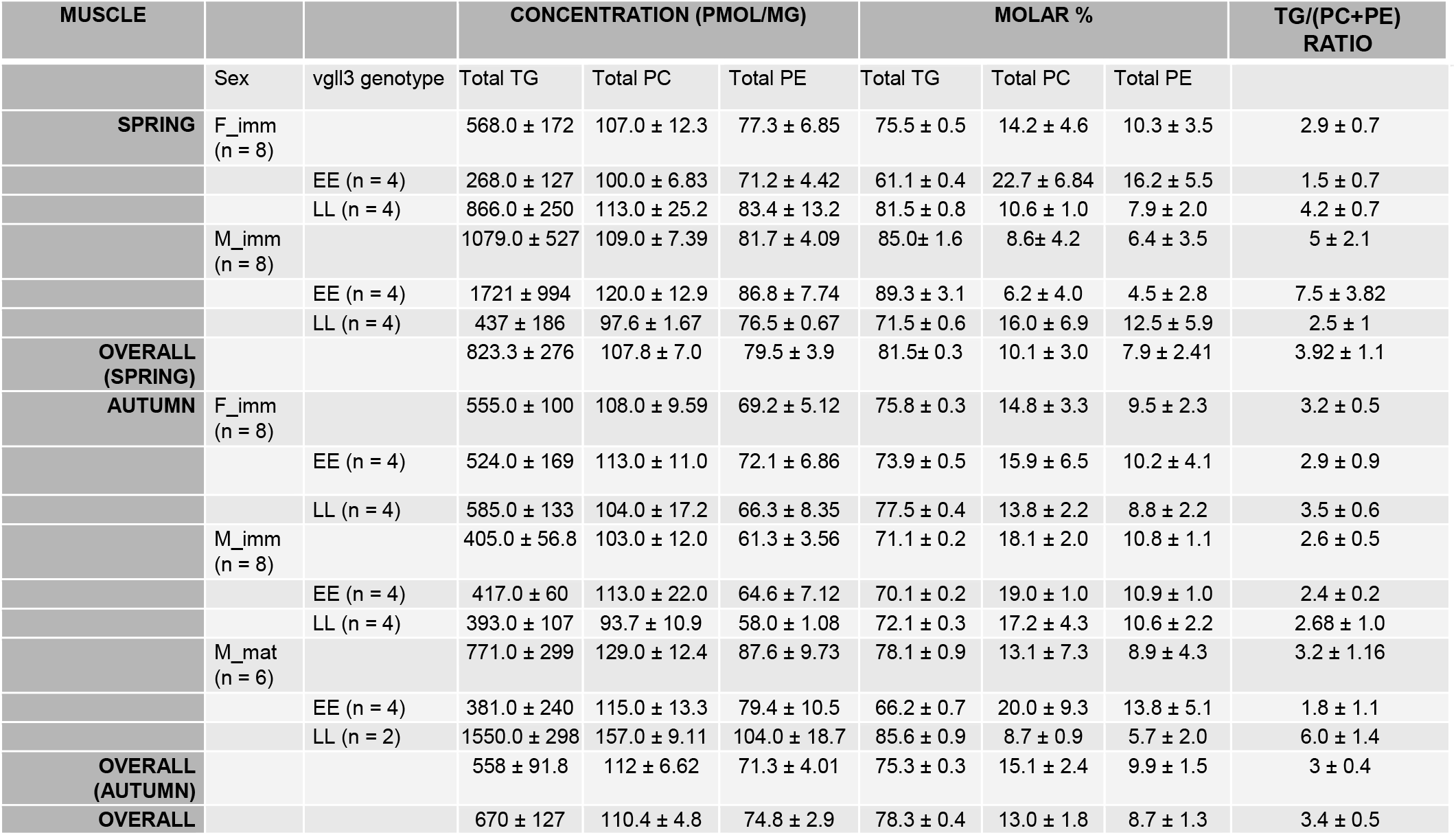
Muscle lipid class concentrations, mol% composition, and TG/(PC +PE) ratios per time point, sex and *vgll3* genotype (EE = early maturation genotype, LL= late maturation genotype) of 2-year old immature (imm) or mature (mat) male (M) and female (F) Atlantic salmon (mean ± SD).

| MUSCLE |  |  | CONCENTRATION (PMOL/MG) |  |  | MOLAR % |  |  | TG/(PC+PE)<br>RATIO |
| --- | --- | --- | --- | --- | --- | --- | --- | --- | --- |
|  | Sex | *vgll3* genotype | Total TG | Total PC | Total PE | Total TG | Total PC | Total PE |  |
| **SPRING** | F_imm<br>(n = 8) | | 568.0 ± 172 | 107.0 ± 12.3 | 77.3 ± 6.85 | 75.5 ± 0.5 | 14.2 ± 4.6 | 10.3 ± 3.5 | 2.9 ± 0.7 |
| | | EE (n = 4) | 268.0 ± 127 | 100.0 ± 6.83 | 71.2 ± 4.42 | 61.1 ± 0.4 | 22.7 ± 6.84 | 16.2 ± 5.5 | 1.5 ± 0.7 |
| | | LL (n = 4) | 866.0 ± 250 | 113.0 ± 25.2 | 83.4 ± 13.2 | 81.5 ± 0.8 | 10.6 ± 1.0 | 7.9 ± 2.0 | 4.2 ± 0.7 |
| | M_imm<br>(n = 8) | | 1079.0 ± 527 | 109.0 ± 7.39 | 81.7 ± 4.09 | 85.0 ± 1.6 | 8.6 ± 4.2 | 6.4 ± 3.5 | 5 ± 2.1 |
| | | EE (n = 4) | 1721 ± 994 | 120.0 ± 12.9 | 86.8 ± 7.74 | 89.3 ± 3.1 | 6.2 ± 4.0 | 4.5 ± 2.8 | 7.5 ± 3.82 |
| | | LL (n = 4) | 437 ± 186 | 97.6 ± 1.67 | 76.5 ± 0.67 | 71.5 ± 0.6 | 16.0 ± 6.9 | 12.5 ± 5.9 | 2.5 ± 1 |
| **OVERALL**<br>**(SPRING)** | | | 823.3 ± 276 | 107.8 ± 7.0 | 79.5 ± 3.9 | 81.5 ± 0.3 | 10.1 ± 3.0 | 7.9 ± 2.41 | 3.92 ± 1.1 |
| **AUTUMN** | F_imm<br>(n = 8) | | 555.0 ± 100 | 108.0 ± 9.59 | 69.2 ± 5.12 | 75.8 ± 0.3 | 14.8 ± 3.3 | 9.5 ± 2.3 | 3.2 ± 0.5 |
| | | EE (n = 4) | 524.0 ± 169 | 113.0 ± 11.0 | 72.1 ± 6.86 | 73.9 ± 0.5 | 15.9 ± 6.5 | 10.2 ± 4.1 | 2.9 ± 0.9 |
| | | LL (n = 4) | 585.0 ± 133 | 104.0 ± 17.2 | 66.3 ± 8.35 | 77.5 ± 0.4 | 13.8 ± 2.2 | 8.8 ± 2.2 | 3.5 ± 0.6 |
| | M_imm<br>(n = 8) | | 405.0 ± 56.8 | 103.0 ± 12.0 | 61.3 ± 3.56 | 71.1 ± 0.2 | 18.1 ± 2.0 | 10.8 ± 1.1 | 2.6 ± 0.5 |
| | | EE (n = 4) | 417.0 ± 60 | 113.0 ± 22.0 | 64.6 ± 7.12 | 70.1 ± 0.2 | 19.0 ± 1.0 | 10.9 ± 1.0 | 2.4 ± 0.2 |
| | | LL (n = 4) | 393.0 ± 107 | 93.7 ± 10.9 | 58.0 ± 1.08 | 72.1 ± 0.3 | 17.2 ± 4.3 | 10.6 ± 2.2 | 2.68 ± 1.0 |
| | M_mat<br>(n = 6) | | 771.0 ± 299 | 129.0 ± 12.4 | 87.6 ± 9.73 | 78.1 ± 0.9 | 13.1 ± 7.3 | 8.9 ± 4.3 | 3.2 ± 1.16 |
| | | EE (n = 4) | 381.0 ± 240 | 115.0 ± 13.3 | 79.4 ± 10.5 | 66.2 ± 0.7 | 20.0 ± 9.3 | 13.8 ± 5.1 | 1.8 ± 1.1 |
| | | LL (n = 2) | 1550.0 ± 298 | 157.0 ± 9.11 | 104.0 ± 18.7 | 85.6 ± 0.9 | 8.7 ± 0.9 | 5.7 ± 2.0 | 6.0 ± 1.4 |
| **OVERALL**<br>**(AUTUMN)** | | | 558 ± 91.8 | 112 ± 6.62 | 71.3 ± 4.01 | 75.3 ± 0.3 | 15.1 ± 2.4 | 9.9 ± 1.5 | 3 ± 0.4 |
| **OVERALL** | | | 670 ± 127 | 110.4 ± 4.8 | 74.8 ± 2.9 | 78.3 ± 0.4 | 13.0 ± 1.8 | 8.7 ± 1.3 | 3.4 ± 0.5 |

**Table 2:**
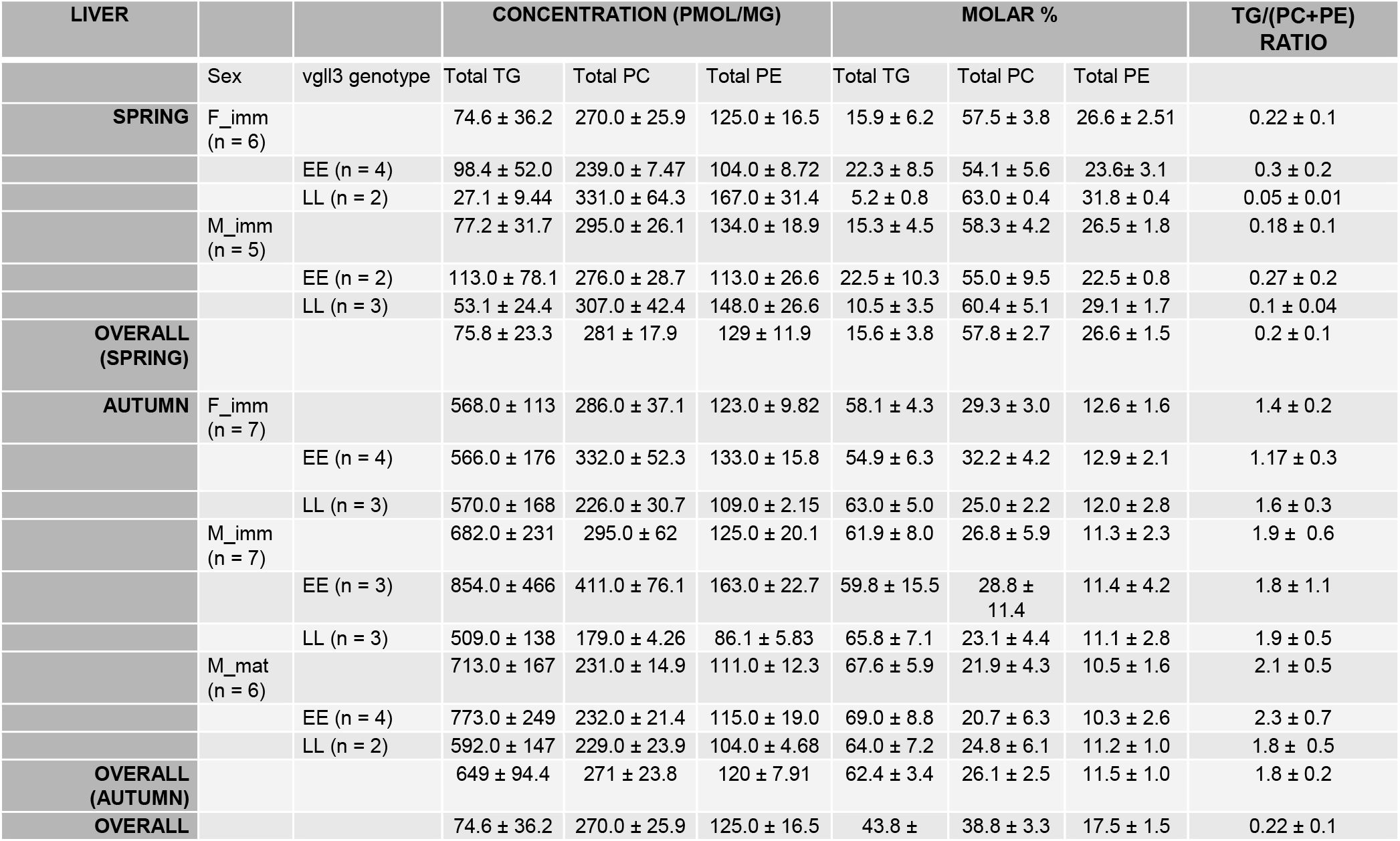
Liver lipid class concentrations, mol% composition, and TG/(PC +PE) ratios per time point, sex and *vgll3* genotype (EE = early maturation genotype, LL= late maturation genotype) of 2-year old immature (imm) or mature (mat) male (M) and female (F) Atlantic salmon (mean ± SD).

| LIVER |  |  | CONCENTRATION (PMOL/MG) |  |  | MOLAR % |  |  | TG/(PC+PE)<br>RATIO |
| --- | --- | --- | --- | --- | --- | --- | --- | --- | --- |
|  | Sex | *vgll3* genotype | Total TG | Total PC | Total PE | Total TG | Total PC | Total PE |  |
| SPRING | F_imm<br>(n = 6) | | 74.6 ± 36.2 | 270.0 ± 25.9 | 125.0 ± 16.5 | 15.9 ± 6.2 | 57.5 ± 3.8 | 26.6 ± 2.51 | 0.22 ± 0.1 |
| | | EE (n = 4) | 98.4 ± 52.0 | 239.0 ± 7.47 | 104.0 ± 8.72 | 22.3 ± 8.5 | 54.1 ± 5.6 | 23.6 ± 3.1 | 0.3 ± 0.2 |
| | | LL (n = 2) | 27.1 ± 9.44 | 331.0 ± 64.3 | 167.0 ± 31.4 | 5.2 ± 0.8 | 63.0 ± 0.4 | 31.8 ± 0.4 | 0.05 ± 0.01 |
| | M_imm<br>(n = 5) | | 77.2 ± 31.7 | 295.0 ± 26.1 | 134.0 ± 18.9 | 15.3 ± 4.5 | 58.3 ± 4.2 | 26.5 ± 1.8 | 0.18 ± 0.1 |
| | | EE (n = 2) | 113.0 ± 78.1 | 276.0 ± 28.7 | 113.0 ± 26.6 | 22.5 ± 10.3 | 55.0 ± 9.5 | 22.5 ± 0.8 | 0.27 ± 0.2 |
| | | LL (n = 3) | 53.1 ± 24.4 | 307.0 ± 42.4 | 148.0 ± 26.6 | 10.5 ± 3.5 | 60.4 ± 5.1 | 29.1 ± 1.7 | 0.1 ± 0.04 |
| OVERALL<br>(SPRING) | | | 75.8 ± 23.3 | 281 ± 17.9 | 129 ± 11.9 | 15.6 ± 3.8 | 57.8 ± 2.7 | 26.6 ± 1.5 | 0.2 ± 0.1 |
| AUTUMN | F_imm<br>(n = 7) | | 568.0 ± 113 | 286.0 ± 37.1 | 123.0 ± 9.82 | 58.1 ± 4.3 | 29.3 ± 3.0 | 12.6 ± 1.6 | 1.4 ± 0.2 |
| | | EE (n = 4) | 566.0 ± 176 | 332.0 ± 52.3 | 133.0 ± 15.8 | 54.9 ± 6.3 | 32.2 ± 4.2 | 12.9 ± 2.1 | 1.17 ± 0.3 |
| | | LL (n = 3) | 570.0 ± 168 | 226.0 ± 30.7 | 109.0 ± 2.15 | 63.0 ± 5.0 | 25.0 ± 2.2 | 12.0 ± 2.8 | 1.6 ± 0.3 |
| | M_imm<br>(n = 7) | | 682.0 ± 231 | 295.0 ± 62 | 125.0 ± 20.1 | 61.9 ± 8.0 | 26.8 ± 5.9 | 11.3 ± 2.3 | 1.9 ± 0.6 |
| | | EE (n = 3) | 854.0 ± 466 | 411.0 ± 76.1 | 163.0 ± 22.7 | 59.8 ± 15.5 | 28.8 ± 11.4 | 11.4 ± 4.2 | 1.8 ± 1.1 |
| | | LL (n = 3) | 509.0 ± 138 | 179.0 ± 4.26 | 86.1 ± 5.83 | 65.8 ± 7.1 | 23.1 ± 4.4 | 11.1 ± 2.8 | 1.9 ± 0.5 |
| | M_mat<br>(n = 6) | | 713.0 ± 167 | 231.0 ± 14.9 | 111.0 ± 12.3 | 67.6 ± 5.9 | 21.9 ± 4.3 | 10.5 ± 1.6 | 2.1 ± 0.5 |
| | | EE (n = 4) | 773.0 ± 249 | 232.0 ± 21.4 | 115.0 ± 19.0 | 69.0 ± 8.8 | 20.7 ± 6.3 | 10.3 ± 2.6 | 2.3 ± 0.7 |
| | | LL (n = 2) | 592.0 ± 147 | 229.0 ± 23.9 | 104.0 ± 4.68 | 64.0 ± 7.2 | 24.8 ± 6.1 | 11.2 ± 1.0 | 1.8 ± 0.5 |
| OVERALL<br>(AUTUMN) | | | 649 ± 94.4 | 271 ± 23.8 | 120 ± 7.91 | 62.4 ± 3.4 | 26.1 ± 2.5 | 11.5 ± 1.0 | 1.8 ± 0.2 |
| OVERALL | | | 74.6 ± 36.2 | 270.0 ± 25.9 | 125.0 ± 16.5 | 43.8 ± | 38.8 ± 3.3 | 17.5 ± 1.5 | 0.22 ± 0.1 |

### Statistical Analysis

Each lipid class or species was log transformed and scaled (each value was subtracted by the mean of the variable, followed by dividing by the standard deviation) before analysis (van den Berg et al., 2006). Exploratory principal component analyses (PCA) were carried out first for each tissue separately using mol% of lipid species to assess the relationship between independent variables. Additionally, only immature females and males were included when testing the differences of total concentration of lipid class between spring and autumn time points and genotypes. Linear mixed effects models were used to test response variable (TG, PC, PE) interactions with fixed effects including Sex (male/female), *vgll3* Genotype (*vgll3*EE/vgll3*\*LL), Time Point (spring/autumn), Maturation Status (immature/mature), and fitting random terms for tank and family. All statistical analyses were conducted using R version 4.2.0 and RStudio 2022.07.2 with packages factoextra, lmertest, lme4, performance for analysis and tidyverse, ggplot2 and ggpubr for visualizing the data and results (Wickham, 2011; Wickham et al., 2019).

## Results

### Muscle lipid profile stability and Liver TG enrichment between seasons

Lipid concentrations of TG, PC and PE in the muscle and liver of juvenile Atlantic salmon are reported in Table 1 and Table 2, respectively. Muscle lipids were composed primarily of the TG lipid class in both the spring (81.5%) and the autumn (75.3%) (Table 1). No significant differences were detected in the concentrations of each lipid class in the muscle between individuals of different *vgll3* genotype, sex or season (Table 3). In contrast, the liver lipid concentrations showed statistically significantly differences for each lipid class studied (Tables 2, 3). PC had the highest mol% in liver lipids in the spring (57.8%) while TG had the highest mol% in liver in the autumn displaying a fourfold enrichment from spring to autumn (15.6 to 62.4%). The liver ratio of TG/(PC+PE) also changed by almost an order of magnitude from the spring to the autumn (0.2 to 1.8).

**Table 3:** Model results based on samples from immature males and females with sex (male/female), *vgll3* genotype (*vgll3*\*EE = early maturation/*vgll3*\*LL = late maturation), and time point (spring/autumn).

| Muscle |  |  |  |  |  |  |
| --- | --- | --- | --- | --- | --- | --- |
| TG | Terms | Estimate | SE | df | t-value | p |
|  | Time_Point | 0.144 | 0.608 | 25.772 | 0.237 | 0.814 |
|  | Sexm | -0.372 | 2.498 | 25.914 | -0.149 | 0.883 |
|  | Vgll3*LL | 0.785 | 2.402 | 24.820 | 0.327 | 0.747 |
|  | Time_Point:Sexm | 0.084 | 0.706 | 25.893 | 0.119 | 0.906 |
|  | Time_Point: Vgll3*LL | -0.289 | 0.679 | 24.819 | -0.425 | 0.675 |
| PC |  | Estimate | SE | df | t-value | p |
|  | Time_Point | -0.068 | 0.168 | 25.989 | -0.401 | 0.692 |
|  | Sexm | -0.388 | 0.684 | 24.029 | -0.567 | 0.576 |
|  | Vgll3*LL | -0.142 | 0.686 | 25.455 | -0.208 | 0.837 |
|  | Time_Point:Sexm | 0.112 | 0.193 | 23.637 | 0.578 | 0.569 |
|  | Time_Point: Vgll3*LL | 0.070 | 0.194 | 25.274 | 0.363 | 0.720 |

| PE |  | Estimate | SE | df | t-value | p |
| --- | --- | --- | --- | --- | --- | --- |
|  | Time_Point | 0.056 | 0.120 | 26.000 | 0.463 | 0.647 |
|  | Sexm | -0.623 | 0.492 | 26.000 | -1.267 | 0.216 |
|  | Vgll3*LL | -0.306 | 0.492 | 26.000 | -0.621 | 0.540 |
|  | Time_Point:Sexm | 0.183 | 0.139 | 26.000 | 1.318 | 0.199 |
|  | Time_Point: Vgll3*LL | 0.100 | 0.139 | 26.000 | 0.721 | 0.478 |
| Liver |  |  |  |  |  |  |
| TG | Terms | Estimate | SE | df | t-value | p |
|  | Time_Point | -1.865 | 0.489 | 17.860 | -3.812 | **0.001** |
|  | Sexm | -0.422 | 2.269 | 17.999 | -0.186 | 0.854 |
|  | Vgll3*LL | 3.886 | 2.253 | 17.884 | 1.725 | 0.102 |
|  | Time_Point:Sexm | 0.060 | 0.632 | 18.000 | 0.094 | 0.926 |
|  | Time_Point: Vgll3*LL | -0.969 | 0.626 | 17.860 | -1.548 | 0.139 |
| PC |  | Estimate | SE | df | t-value | p |
|  | Time_Point | -0.373 | 0.142 | 17.186 | -2.634 | **0.017** |
|  | Sexm | -0.263 | 0.674 | 17.690 | -0.391 | 0.701 |
|  | Vgll3*LL | -2.601 | 0.639 | 15.903 | -4.070 | **0.001** |
|  | Time_Point:Sexm | 0.068 | 0.187 | 17.581 | 0.362 | 0.721 |
|  | Time_Point: Vgll3*LL | 0.798 | 0.176 | 15.310 | 4.520 | **0.000** |
| PE |  | Estimate | SE | df | t-value | p |
|  | Time_Point | -0.286 | 0.145 | 17.682 | -1.965 | 0.065 |
|  | Sexm | 0.060 | 0.689 | 17.927 | 0.086 | 0.932 |
|  | Vgll3*LL | -2.595 | 0.662 | 16.265 | -3.922 | **0.001** |
|  | Time_Point:Sexm | -0.014 | 0.191 | 17.873 | -0.072 | 0.944 |
|  | Time_Point: Vgll3*LL | 0.748 | 0.183 | 15.799 | 4.085 | **0.001** |

### Clear lipid species profile differences between seasons

In muscle tissue, 58, 32, and 14 lipid species were detected for TG, PC, and PE, respectively, while in the liver, 44, 39, and 15 lipid species were detected for TG, PC, and PE, respectively. PCA visualization of individual-level lipid species patterns revealed a clear separation in the lipid species composition between seasons in both tissues (Figure 2). Specifically, the spring and autumn samples of muscle separate along the second principal component explaining 21% of the variation (Figure 2A). In muscle, the degree of unsaturation of TG species was in general higher in the spring samples (total of 6-8 double bonds in the acyl chains of the molecule) than in the autumn samples (3-5 double bonds) (Figure 2A). In contrast, the spring and autumn samples of liver separate along the first component explaining 89% of the variation (Figure 2B), likely due to the dramatic enrichment in TG mol% values from spring to autumn. In both tissues, the quantitatively most important PC and PE species 38:6 is enriched in the spring samples. The PCA biplot of muscle shows an interesting shift, since in the spring the lipid species that have the highest double bond contents (more biochemically available, (Raclot, 2003)) are TG species (e.g., TGs 56:7 and 56:8) and in the autumn the species with the highest double bond contents are phospholipid species (PC 40:7 and PE 40:7, most likely containing a 22:6n-3 acyl chain).

**Figure 2:**
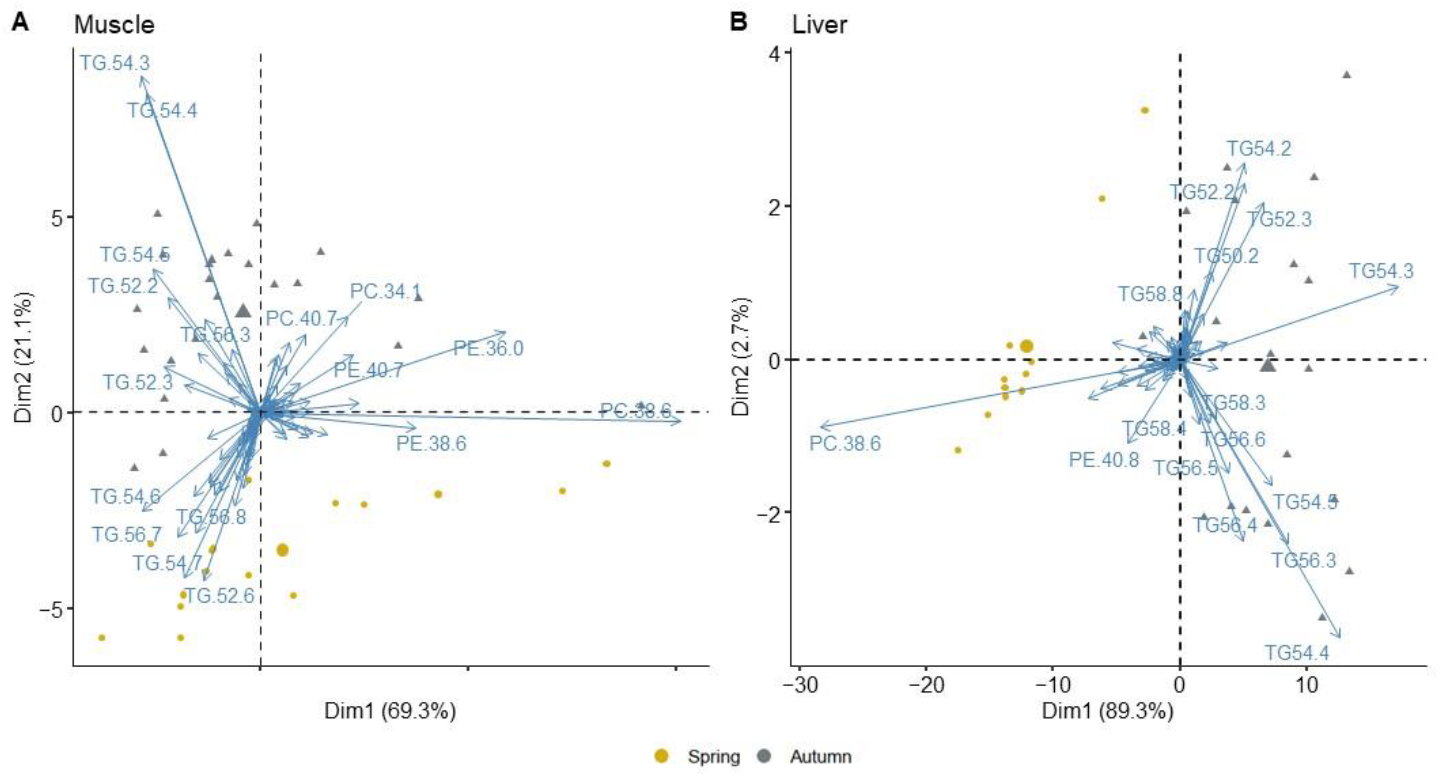
PCA showing principal component 1 and 2 for mol% of lipid species in A) muscle (n = 38), and B) liver (n = 30). Yellow dots indicate spring and gray triangles indicate autumn samples with the largest symbols of each representing the mean of each season group. Arrows show loadings of separate lipid species contributing most to each principal component.

### Vgll3 and season effects on muscle and liver lipid concentrations

There were no statistically significant differences of lipid class concentrations in the muscle when comparing spring and autumn samples, nor when comparing sexes or *vgll3* genotypes (Table 3, Figure 3). In contrast, in liver tissue, several statistically significant differences in lipid class concentrations were identified (Table 3). In the liver, TG concentrations were significantly higher in the autumn than in the spring (Table 3, Figure 4A). There were also differences in PC and PE concentrations between *vgll3* genotypes and spring and autumn: immature *vgll3*\*EE individuals increase or maintain their liver PC and PE concentrations from spring to autumn, whereas these concentrations decrease in *vgll3*LL* individuals towards the autumn (Table 3, Figure 4B, C). This resulted in a highly statistically significant interaction between *vgll3* genotype and season for both PC and PE concentrations, whereby liver PC and PE concentrations of *vgll3*\*LL individuals decreased from spring to autumn while the seasonal change in concentration was the opposite in *vgll3*\*EE individuals (Table 3, Figure 4B, C).

**Figure 3:**
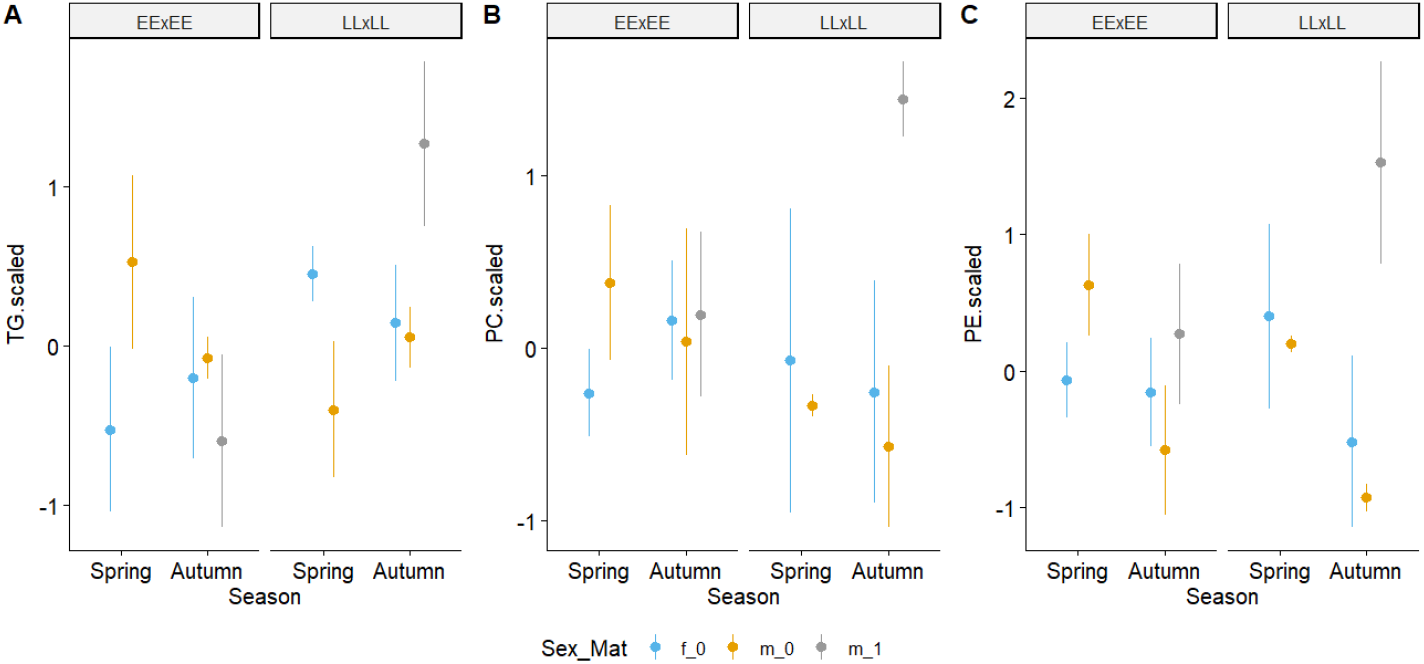
Scaled A) triacylglycerol (TG), B) phosphatidylcholine (PC) and C) phosphatidylethanolamine (PE) concentrations of muscle in vgll3*EE (EExEE) and vgll3*LL (LLxLL) immature females (f_0), immature males (m_0), and mature males (m_1) between the spring and autumn.

**Figure 4:**
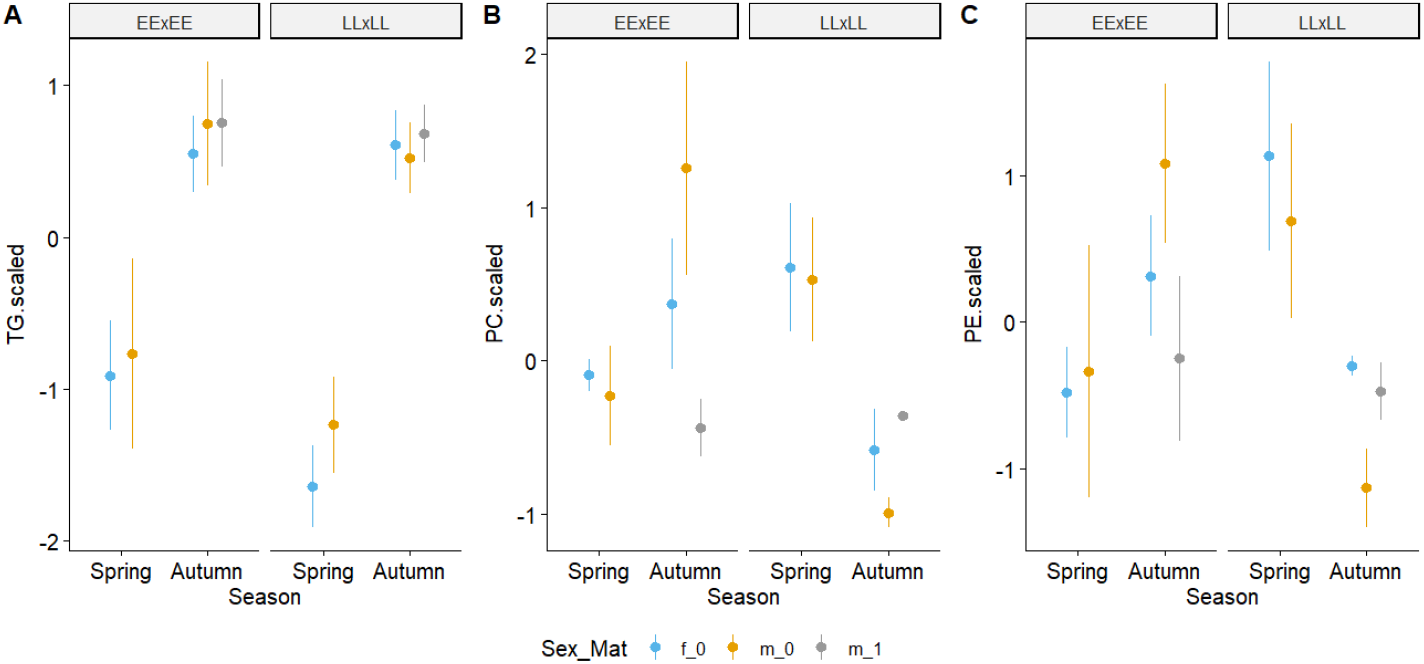
Scaled A) triacylglycerol (TG), B) phosphatidylcholine (PC) and C) phosphatidylethanolamine (PE) concentrations of liver in vgll3*EE (EExEE) and vgll3*LL (LLxLL) immature females (f_0), immature males (m_0), and mature males (m_1) between the spring and autumn.

### Maturation, sex, and vgll3

Mature males with the *vgll3*\*LL genotype had significantly higher TG and PC concentrations in the muscle than *vgll3*\*EE individuals (Figure 3 A & B, Table 4). However, this was based on data from only two *vgll3*\*LL mature males. No differences in lipid concentrations were observed between immature males and females.

**Table 4:** Model results including mature males to test for lipid class and *vgll3* association (Status (male/female: immature/mature) *vgll3* Genotype (*vgll3*\*EE = early maturation /*vgll3*\*LL = late maturation).

| Muscle |  |  |  |  |  |  |
| --- | --- | --- | --- | --- | --- | --- |
| TG | Terms | Estimate | SE | df | t-value | p |
|  | Sex_Mat | -0.849 | 0.573 | 17.893 | -1.482 | 0.156 |
|  | Vgll3*LL | -0.262 | 0.522 | 16.494 | -0.502 | 0.623 |
|  | Sex_Mat:Vgll3*LL | 2.402 | 0.939 | 17.960 | 2.560 | 0.020 |

| PC |  |  |  |  |  |  |
| --- | --- | --- | --- | --- | --- | --- |
|  |  | Estimate | SE | df | t-value | p |
|  | Sex_Mat | 0.076 | 0.466 | 11.574 | 0.164 | 0.873 |
|  | Vgll3*LL | -0.598 | 0.485 | 9.531 | -1.232 | 0.248 |
|  | Sex_Mat: Vgll3*LL | 1.787 | 0.792 | 13.550 | 2.256 | 0.041 |
| PE |  | Estimate | SE | df | t-value | p |
|  | Sex_Mat | 0.730 | 0.478 | 6.217 | 1.527 | 0.176 |
|  | Vgll3*LL | -0.004 | 0.567 | 8.449 | -0.007 | 0.995 |
|  | Sex_Mat:Vgll3*LL | 1.060 | 0.842 | 9.658 | 1.258 | 0.238 |
| Liver |  |  |  |  |  |  |
| TG |  | Estimate | SE | df | t-value | p |
|  | Sex_Mat | 0.610 | 0.622 | 13.000 | 0.980 | 0.345 |
|  | Vgll3*LL | -0.499 | 0.562 | 13.000 | -0.888 | 0.391 |
|  | Sex_Mat: Vgll3*LL | 0.425 | 0.981 | 13.000 | 0.434 | 0.672 |
| PC |  | Estimate | SE | df | t-value | p |
|  | Sex_Mat | -1.3931 | 0.6545 | 13 | -2.128 | 0.053 |
|  | Vgll3*LL | -1.3468 | 0.5908 | 13 | -2.279 | 0.0402 |
|  | Sex_Mat: Vgll3*LL | 1.3253 | 1.0311 | 13 | 1.285 | 0.2211 |
| PE |  | Estimate | SE | df | t-value | p |
|  | Sex_Mat | -0.801 | 0.784 | 13.000 | -1.023 | 0.325 |
|  | Vgll3*LL | -0.783 | 0.707 | 13.000 | -1.107 | 0.288 |
|  | Sex_Mat: Vgll3*LL | 0.554 | 1.234 | 13.000 | 0.448 | 0.661 |

## Discussion

We compared the lipid class concentrations and species profiles of juvenile Atlantic salmon at two key seasonal time points for early salmon life-history: spring and autumn. Lipid levels in the spring likely provide indications of energy use efficiency over the winter, as well as indicate the basal lipid reserves to build on for possible maturation in the coming autumn. However, lipid levels in the autumn reflect resources accumulated over the summer, and both the initial and newly acquired lipids potentially contribute to the available total energy source for the spawning event in the autumn (Rowe et al., 1991). Initiation of early maturation (as a two-year old parr) likely requires physiological changes due to the rapid developmental changes, including gonad development, needed to reproduce (Aksnes et al., 1986). We found that liver TG concentrations increased from spring to autumn, indicating that the liver is potentially an important lipid storage location in Atlantic salmon, as suggested for Arctic charr (Jobling et al., 1998) and white seabream (Cejas et al., 2004). High lipid content in the liver can suggest physiological issues resembling non-alcoholic fatty liver in humans and involving lipid peroxidation and oxidative stress. However, we are currently just beginning to understand these mechanisms in salmon (Espe et al., 2019; Keinänen et al., 2022).

Additionally, we did not detect any significant differences between the sexes. In contrast, a previous study, House et al., (2021), suggested that males had higher concentrations of several lipid classes in the muscle compared to females, but that study was conducted on one year old juveniles which can potentially explain this difference. Another possible explanation for these differing results is that the current study included both mature and immature males whereas the previous study was conducted on younger and solely immature males, but may have included males with higher lipid levels that may have matured the following year. A more detailed study of male vs. female lipid differences across a longer time period would be useful for better understanding sex-specific lipid allocation strategies.

Despite muscle lipid class concentrations not showing any marked changes from spring to autumn, the individual lipid species composition changed markedly (Fig. 2A). The muscle TG species in the spring were highly unsaturated species that are used much less for fatty acid β-oxidation, potentially indicating acclimatization to low water temperature (Colombo et al., 2022; Wang et al., 2021). In contrast, the TG species accumulated during summer and detected in the autumn samples had a lower degree of unsaturation, which, during higher summer and autumn water temperature and physiological activity, may protect juvenile salmon against oxidative stress (Gray, 1978; Kjær et al., 2008). Adult female salmon with the largest reserves of polyunsaturated lipid are known to suffer from oxidative stress during spawning and fail to breed (Keinänen et al., 2022; Vuorinen et al., 2020). Therefore, the lipid species profiles of the juvenile salmon of this study suggest that distributing highly unsaturated fatty acids into phospholipids may reduce the rates of polyunsaturated fatty acid peroxidation and thereby reduce oxidative stress..

Perhaps the most noteworthy finding of this study is the stark contrast in the direction of membrane lipid (PC and PE) concentration changes between seasons in *vgll3*\*EE individuals compared to *vgll3*\*LL individuals. This can possibly be explained by two scenarios, one involving genotype-specific differences in lipid synthesis, and another involving genotype-specific differences in lipid storage mechanisms. In the first scenario, the observed lipid concentrations could reflect differences in endoplasmic reticulum (ER) volume in differing *vgll3* genotype individuals in the liver. The higher membrane lipid concentrations increasing from spring to autumn in *vgll3*\*EE individuals could be maintaining a more stable capacity of ER functions compared to the *vgll3*\*LL individuals decreasing from spring to autumn. The ER is one of the major sites of protein synthesis and fatty acid and lipid metabolism, including *de novo* synthesis of phospholipids and TG, while it also produces lipoprotein particles for transport of diverse biomolecules throughout the body, the TGs largely carried to the main storage sites, such as muscle myosepta and visceral adipose tissue (Jensen-Urstad & Semenkovich, 2012; Alves-Bezerra & Cohen, 2017). Thus, this increase of membrane lipid concentrations in *vgll3*\*EE individuals may increase or retain capability for ER functions across seasons compared to the decrease of membrane lipids seen in *vgll3*\*LL individuals. For the second scenario, individuals could be storing lipid droplets in liver and other tissues in a genotypic-specific manner, resulting in the estimated contrast in the direction of membrane lipid (PC and PE) concentration changes from spring to autumn in *vgll3*\*EE individuals compared to *vgll3*\*LL individuals. Specifically, lipid droplets stored in the cytoplasm are the main storage organelles for metabolic energy in most cells (Prévost et al., 2018). The membrane lipid (PC and PE) concentration decrease from spring to autumn in *vgll3*\*LL individuals could be due to them storing larger lipid droplets in the liver compared to *vgll3*\*EE individuals, and thus increasing the mass of storage lipid at the expense of ER network mass. However, a more complete understanding of these genotype-based differences requires additional research. For example, *vgll3* genotype specific differences in transcriptomic data identified lipid metabolism genes, such as fatty acyl desaturases (FADS) and elongases of very long chain fatty acids (ELOVLs), known to be involved in fatty acid structural modifications (Datsomor et al., 2022; Kabeya et al., 2018) and also *vgll3* genotype specific differences in histological patterns of lipid droplet determination (hyperplasic vs hypertrophy) (Caballero et al., 2002).

Our findings in juvenile Atlantic salmon provide first hints at mechanisms by which *vgll3* contributes to the maintenance of salmon lipid reserves and metabolic capability across seasons. If *vgll3*\*EE individuals do indeed have increased ER metabolic activity with an increased concentration of membrane lipids from spring to autumn compared to the opposite trend in *vgll3*LL* individuals, it would imply they may have a higher capacity for protein synthesis and thereby a better capability for production of phospholipid and TG across seasons. It would explain why *vgll3*\*EE individuals maintained a higher body condition in the spring (House et al. 2023), which increases the probability of the ability to mature in the autumn. These results are in line with previous studies that reported for *vgll3*\*EE relative to *vgll3*\*LL individuals a higher aerobic scope (Prokkola et al., 2022), body condition (Debes et al., 2021) and also a more seasonally stable body condition (House et al., 2023).

In conclusion, we found that seasonality has a major impact on Atlantic salmon lipid profiles and with *vgll3* specific effects on membrane lipid concentrations. The precise mechanism linking *vgll3* with lipid metabolism and storage is still not clear, but this study adds to the increasing indirect evidence supporting the notion that such a mechanism exists. Atlantic salmon juveniles seem to exhibit genotype specific lipid profiles with *vgll3*\*EE individuals increasing membrane lipid concentrations between spring and autumn while *vgll3*\*LL individuals doing the opposite. Future work investigating *vgll3* specific genotype differences in the expression of lipid-metabolism related genes would help understand the mechanism by which *vgll3* influences maturation timing. Further, assessment of lipid profiles in additional tissues and at additional time points, ideally over a longer time period, would allow for a more systematic assessment of the processes influencing early maturation probability and seasonality effects in juvenile Atlantic salmon.

## Acknowledgements

We acknowledge Ksenia Zueva, Marion Sinclair-Waters, Nico Lorenzen, Spiros Papakostas, and Victoria Pritchard for their assistance in getting fish gametes in Laukaa hatchery; Suvi Ikonen, Anna Toikkanen, Shadi Jansouz, Dorian Jagusch, Andres Salgado, Fin Morrison, Ike Van Gestel, Paul Bangura, and Petra Lijeström for their help in Lammi with husbandry and fish sampling and Outi Ovaskainen; Miakel Kyriacou, Oona Mehtälä, Oliver Andersson, Pirta Palola, Valeria Valanne, and Jacqueline Moustakas-Verho for their help during sampling events; Annukka Ruokolainen, Noora Parre, Shadi Jansouz and Seija Tillanen for their help in the genetics lab and Eirik Åsheim for his help with manuscript discussions and data wrangling, and Tutku Aykanat for help with data analysis and discussions.

